# SDip: A novel graph-based approach to haplotype-aware assembly based structural variant calling in targeted segmental duplications sequencing

**DOI:** 10.1101/2020.02.25.964445

**Authors:** David Heller, Martin Vingron, George Church, Heng Li, Shilpa Garg

## Abstract

Segmental duplications are important for understanding human diseases and evolution. The challenge to distinguish allelic and duplication sequences has hindered their phased assembly as well as characterization of structural variant calls. Here we have developed a novel graph-based approach that leverages single nucleotide differences in overlapping reads to distinguish allelic and duplication sequences information from long read accurate PacBio HiFi sequencing. These differences enable to generate allelic and duplication-specific overlaps in the graph to spell out phased assembly used for structural variant calling. We have applied our method to three public genomes: CHM13, NA12878 and HG002. Our method resolved 86% of duplicated regions fully with contig N50 up to 79 kb and produced <800 structural variant phased calls, outperforming state-of-the-part SDA method in terms of all metrics. Furthermore, we demonstrate the importance of phased assemblies and variant calls to the biologically-relevant duplicated genes such as SMN1, SRGAP2C, NPY4R and FAM72A. Our phased assemblies and accurate variant calling specifically in duplicated regions will enable the study of the evolution and adaptation of various species.

## Introduction

Segmental duplications (SDs) are special types of repeats of size >1 Kbp that are repeated within the genome with at least 90% sequence identity in either tandem or interspersed organization ^1,2^. They are a source of phenotypic variability in disease and health. SDs are among the most important sources of evolution, a common cause of genomic structural variation and several are associated with diseases of genomic origin, including schizophrenia and autism ^3–6^.

In recent years, single-molecule sequencing approaches such as PacBio SMRT sequencing or Oxford Nanopore sequencing ^7^ have enabled the de-novo assembly of genomes to a considerably higher quality than was possible with traditional short-read methods. Both technologies produce longer reads several kbps in length which allows to resolve many repeats and complex genomic regions because long enough reads can encompass whole expanded repeats, and can be anchored using the flanking unique sequences ^8^. However, both technologies come at the cost of a relatively high error rate of 10-15%. This causes problems in regions of SDs where different paralogues and alleles can be highly similar and impossible to distinguish with error-prone reads ^9,10^. As a consequence, copies of duplicated regions are collapsed in most assembly approaches. The availability of consensus reads, such as PacBio HiFi reads ^11^, with an accuracy of ~99% accuracy and average length of ~15kb, that are derived from multiple sequencing rounds of the same fragment, offers enormous opportunities to potentially resolve duplicated regions with high repeat identities. There is a pressing need for bioinformatics methods designed specifically to take advantage of combined characteristics of accuracy and longer length from HiFi sequencing to better resolve duplication regions with high repeat identities.

There are only a few tools such as Falcon Unzip ^12,13^ and TrioCanu ^14^ that can generate phased contigs in highly divergent repeats, but they both have limitations in more similar regions such as SDs. The high similarity between different paralogues and alleles of an SD causes many of these duplicated regions to be collapsed ^9,10^). For their tool SDA, Vollger et al. used long-read sequencing to partition paralogues based on paralogous sequence variants (PSVs) into uncollapsed contigs. However, SDA has the limitation to produce diploid contigs and is not specifically designed for HiFi data. Currently, there are no tools available that can generate accurate phased contigs in SDs and other complex repeats such as LINEs and SINEs.

Here we propose our novel graph-based method SDip that leverages information from long accurate PacBio HiFi reads to distinguish reads from different haplotypes and paralogues in the process of generating an overlap graph thus generating an haplotype and repeat-aware graph. Specifically, we consider single nucleotide variations (SNVs) to distinguish reads belonging to different haplotypes and paralogues from overlapping reads. Therefore, only reads belonging to the same haplotype and paralogue are connected in the overlap graph. Through this graph, we thread ultralong nanopore reads that help to spell out phased contigs. After polishing the contigs, we align them to the reference genome to perform calling of structural variants (SVs). SDip, for the first time, has the potential to generate diploid assemblies in complex regions, including in SDs. It can be applied to repeats and SDs over any genome but is specifically designed for PacBio HiFi data. Our method also allows to integrate ultralong nanopore reads although this is optional. Finally, we show that phased contigs from our method can be used for haplotype-aware SV calling and genotyping in complex duplicated regions.

Previous SV genotyping methods for repetitive genomic regions such as PacmonSTR ^15^ and RepeatHMM ^16^ align reads to a reference genome, and then perform sophisticated probability-based comparisons of these reads to the sequence of the repeating unit in the tandem repeat region. More general approaches such as Sniffles ^16,17^, SMRTSV2 ^18^ and SVIM ^19^ call variants from long read alignments or local de-novo assemblies, but have limitations in complex regions, such as segmental duplications. As part of SDip, we extended our previous SV calling method SVIM to leverage information from the alignment of diploid contigs from our method to the reference genome. Thus, SDip produced, for the first time, SV call sets in complex segmental duplications that have potential applications in evolutionary and comparative genomics.

We apply SDip to SDs over three human genome assemblies derived from haploid (CHM13) and diploid genomes (NA12878 and HG002). We reconstruct diploid assemblies through duplicated regions and use them to detect SVs in these regions. We demonstrate that we can produce diploid contigs with an N50 of up to 79 kb and a QV of more than 28 within 24 hours. In all three genomes, we can resolve more than 86% of collapsed regions. Our SV call set comparison to BAC clones is higher compared to state-of-the-art tools. This highlights that this method is accurate and computationally tractable, and its use can be extended to resolve collapsed repeat content and phase in de novo assemblies of other mammalian genomes.

## Results

At the outset of our study, we aligned HiFi data to an assembly of the same individual (base assembly, see Methods) and detected collapsed regions using SDA ^9,10^). For NA12878, we found 486 collapsed regions with an elevated read coverage and an average length of 48.8 kb. When we mapped those regions (23.7 Mbps in total) back to the reference genome (GRCh38), they spanned 40.9 Mbps in sequence and 89.3% (40.1 Mbps) overlapped annotated SD regions. To uncollapse and assemble the collapsed regions, our method SDip leverages key advantages from two data types: the low error-rates of PacBio HiFi data and the longer read lengths from ultralong nanopore reads. By combining both data types, we are able to distinguish both paralogous and allelic sequences in duplicated regions.

We align each collapsed region to the base assembly allowing secondary alignments to find similar regions and then recruit PacBio HiFi reads both from the collapsed and all similar regions (Figure 1). Please note that collapsed regions from the base assembly is one approach for selecting duplicated regions, it depends on the user to use the reference genome or assembly, or alternatively directly use sequencing reads from targeted duplication sequencing. From the recruited reads, our method identifies high-confidence allelic and paralogous single nucleotide variations (SNVs) to generate a haplotype and repeat-aware overlap graph. In this graph, only reads belonging to the same haplotype and paralogue are connected. Subsequently, a series of cleaning and simplification steps are applied to prune tips, simplify small bubbles and remove cross-linking paths in the graph. This process generates a chain of bubbles that can be either simple (representing resolved haplotypes and paralogous) or nested from unresolved cases. Next, ultralong nanopore reads are aligned to this simplified graph to find paths between unique sequences flanking the duplications and to help prevent misassemblies. Finally, a minimum path cover notion is followed to spell out the remaining haplotype paths (including paralogues in nested bubbles) in the graph.

**Fig. 1.**
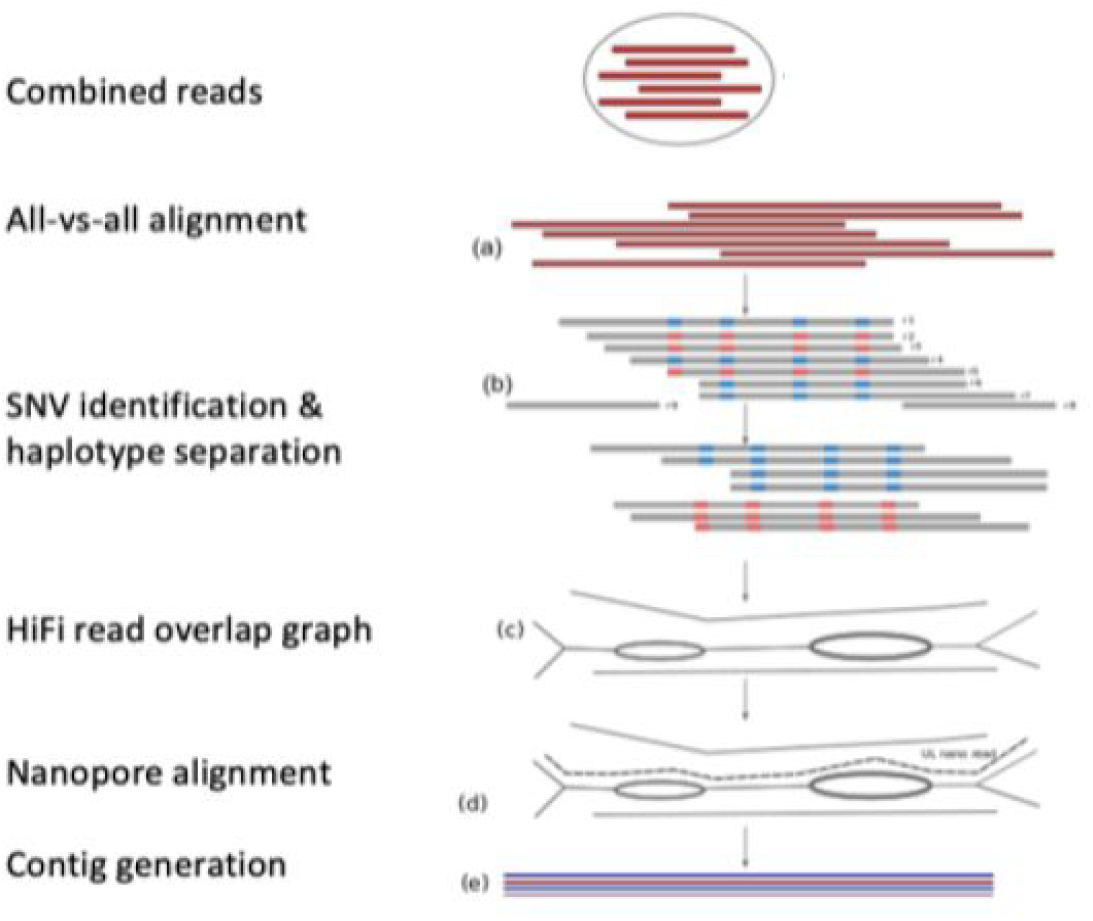
SDip workflow. On combined readsets from similar regions, apply the following steps: a) all-vs-all alignment of reads, b) detect single nucleotide differences (SNVs) in pileup for every target read, then find similar reads to target read potentially belong to same haplotype and repeat using SNP information, c) generate allelic-aware and repeat-aware graph, d) thread ultralong nanopore reads (UL) through graph and e) spell out phased contigs using UL reads and minimum path cover notion.

We applied our method to three human genomes of haploid and diploid nature (see Methods) and validated the results and accuracy based on targeted BAC sequencing (see Methods) and analyses of specific duplicated loci. For NA12878, we were able to assemble 2677 contigs (mean of 5.5 contigs per collapsed region) representing different previously collapsed paralogues and alleles (Table 1). In total, we could resolve 159.3 Mbps of diploid sequence that used to be collapsed in the base assembly. The majority of contigs could be matched as the two chromosomal haplotypes of the same paralogue yielding 55.8 and 55.9 Mbps from either chromosomal haplotype (diploid contigs). An additional 47.6 Mbps remained unmatched and most likely represent paralogues with highly divergent chromosomal haplotypes or paralogues whose chromosomal haplotypes could not be uncollapsed. The mean length of the diploid contigs was 60.5 and 60.6 kbps, respectively for the two chromosomal haplotypes, and the longest contig had a size of 477.2 kbps. Of the 486 regions, 17 failed to generate an assembly, mainly because of cycles in the overlap graph. Another 22 regions were excluded from the analysis because of a high number of recruited reads or similar regions resulting in too many read overlaps.

**Table 1.** Assembly statistics. N50: 50% of bases are contained in reads/contigs longer than this number. Fully assembled regions: regions spanned end-to-end by at least one contig. Covering contigs per region: contigs spanning a region end-to-end. Misassemblies: prematurely ending BAC-to-contig alignments indicating a contig misassembly. Average QV to BACs: base-level accuracy in Phred scale. Note that no BACs were available for HG002.

| Sample | HG002 (NA24385) |  | NA12878 |  | CHM13 |  |
| --- | --- | --- | --- | --- | --- | --- |
| HiFi read coverage | 29.7 |  | 30.1 |  | 25.2 |  |
| HiFi read N50 | 13,480 |  | 10,004 |  | 10,859 |  |
| Nanopore read coverage | 63.4 |  | 41.0 |  | 122.4 |  |
| Nanopore read coverage (reads>100kb) | 18.1 |  | 3.3 |  | 32.7 |  |
| Assembly algorithm | SDA | SDip | SDA | SDip | SDA | SDip |
| Collapsed regions | 561 |  | 486 |  | 382 |  |
| #contigs / sequence (Mbp) | 1592 / 65.4 | 2672 / 155.0 | 1664 / 62.5 | 2677 / 159.3 | 530 / 20.00 | 1020 / 49.9 |
| Paternal / maternal contig size (Mbp) | - | 53.4 / 53.4 | - | 55.8 / 55.9 |  |  |
| Haploid contig N50 (kbp) | 46.6 | - | 44.5 | - | 39.4 | 51.2 |
| Paternal / maternal contig size N50 (kbp) | - | 62.6 / 61.9 | - | 78.6 / 79.0 |  |  |
| Fully assembled regions | 267 | 458 | 185 | 419 | 86 | 325 |
| Mean/Median covering contigs per region | 0.8 / 0 | 2.8/3 | 0.6 / 0 | 2.8 / 3 | 0.3 / 0 | 1.7 / 2 |
| Misassemblies vs. BACs | - | - | 8 | 11 | 8 | 12 |
| Average QV to BACs | - | - | 24.8 | 28.8 | 24.9 | 23.9 |

When we compared our results to the haploid contigs produced by SDA (Table 1), we found that SDip is able to assemble more than twice as many base pairs (159.3 compared to 62.5 Mbps) and can produce contigs that are considerably more contiguous (diploid contig N50 of 78.6 and 79.0 kbps compared to 44.5 kbps). By aligning our contigs to the reference genome, we found that SDip is also able to resolve substantially more collapsed regions than SDA (419 compared to 185). For each of the resolved regions at least one contig could be assembled that covers the entire region end-to-end while the mean number of spanning contigs per region was 2.8 (0.6 for SDA).

We also analyzed annotated SD regions in hg38 to compare duplications unresolved by the base assembly with those resolved by SDip. To this end, we aligned both the NA12878 base assembly and SDip contigs to the reference genome. 30.9% of SD regions spanning 93.8 Mbps remained unresolved by the base assembly, i.e. they failed to be covered end-to-end by the aligned assembly. SDip was able to resolve 22.4% (11.9 Mbps) of those regions or 15.8% of all SD regions (24.1 Mbps). SDip was able to resolve duplications between 1.0 and 258.4 kbps while regions unresolved by the base assembly had a length between 0.4 and 770.4 kbps. Generally, however, SD regions resolved by SDip tended to have a similar length and percent identity as the regions unresolved by the base assembly (Figure 2a).

**Fig. 2.**
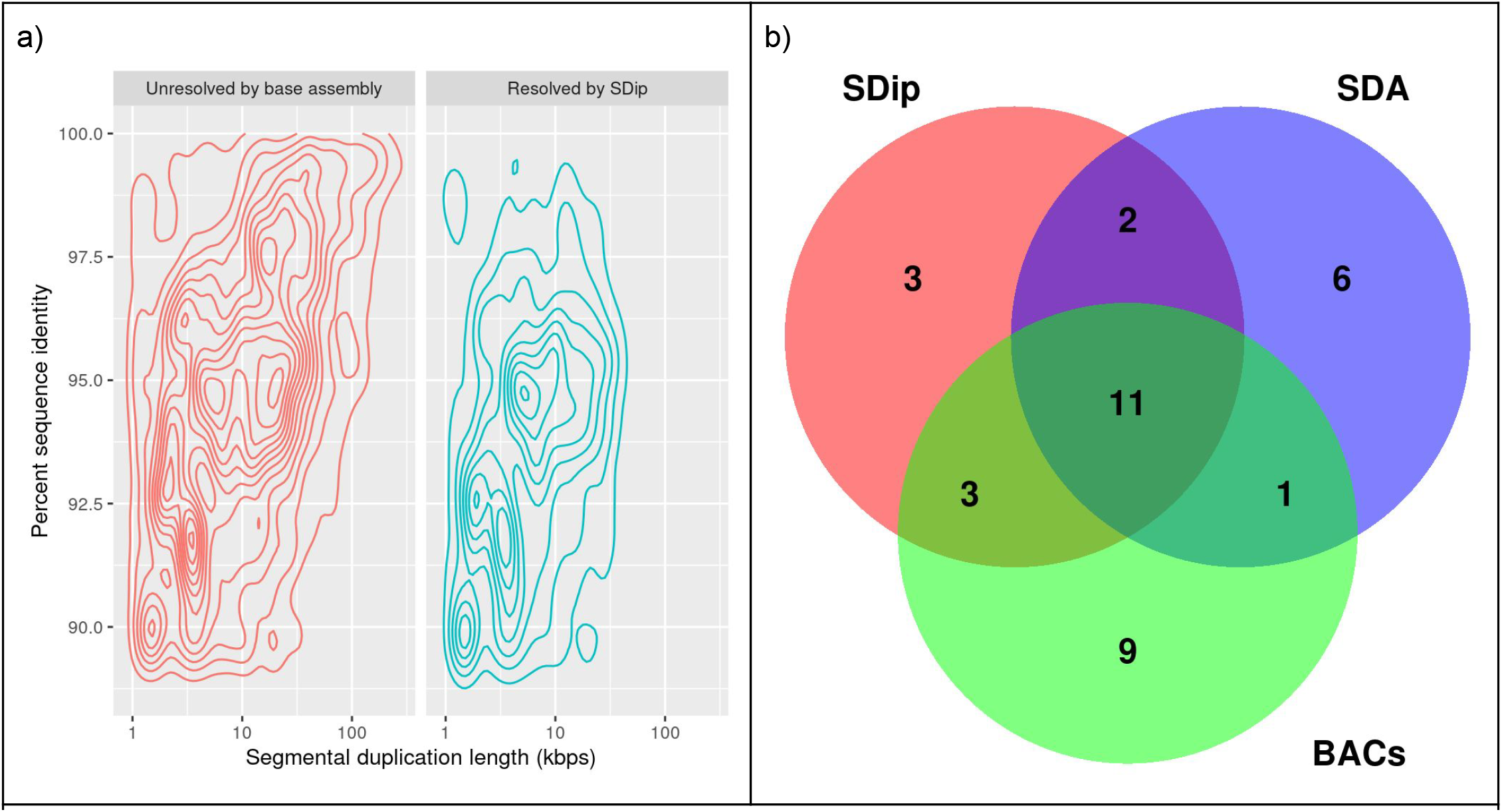
SDip results for the human NA12878 assembly. a) Contour plot of annotated SDs plotted by length and percent sequence identity. Shown are SDs unresolved by the NA12878 base assembly (left) and SDs resolved by SDip (right). b) Venn diagram of three SV callsets from SDip contigs (red), SDA contigs (blue) and BACs (green). All calls were made by SVIM in regions of overlap between SDip contigs, SDA contigs and BACs.

To further investigate the quality of our diploid contigs, we computed quality values from alignments of BAC sequences to the contigs (see Methods). For NA12878, SDip contigs reached a mean QV of 28.8 compared to a mean QV of 24.8 for SDA. From the alignment of BACs we also estimated the number of misassemblies by analyzing alignments that end prematurely, i.e. distant from the ends of both the BAC and the contig. We estimated that 11 contigs from SDip and 8 contigs from SDA have been misassembled.

Diploid contigs assembled by SDip can be aligned to the human reference genome and used for variant calling in complex genomic regions. To this end, we extended our SV caller SVIM to detect, genotype and phase SVs from the alignment of a diploid assembly to the reference genome (GRCh38) ^19^. When we applied SVIM to our contigs for NA12878, we detected 302 deletions and 416 insertions up to 6.1 kbps (deletions) and 4.8 kbps (insertions) in size. 101 / 191 deletions and 193 / 223 insertions were heterozygous or homozygous respectively. We compared SVs detected in our contigs with SVs detected in the BACs and in SDA contigs. Because BACs and contigs do not cover the same genomic regions, we restricted our comparison to regions where alignments of SDip contigs, SDA contigs and BACs overlapped (3.9 Mbps in total). In those regions a total of 35 SVs were detected (Figure 2b). 11 of those were detected in all three datasets while 3, 6 and 9 were detected exclusively from SDip contigs, SDA contigs and BACs, respectively. In general, 16 out of 19 SDip calls could be confirmed by either BACs (14) or SDA contigs (13).

Human-specific segmental duplication genes such as SMN1, SRGAP2C, NPY4R and FAM72A are highly identical (percent identity ~99%) and known to be highly copy number polymorphic^20^. Our analysis of SMN1 and SRGAP2C showed that its genomic locus was collapsed in the base assembly of NA12878 and could be successfully resolved by SDip recreating the sequence and gene models present in the reference genome (Figure 3a, b). For the first time, SDip could produce diploid contigs of SMN1. In the other three genes, SDip produced cleaner fully resolved phased contigs as shown whereas SDA produced only haploid SD sequences. In total, we fully resolved 10 out of 24 known duplicated genes and partly resolved another 7 genes (Supplementary Table 1). SDA, in comparison, was able to resolve only 5 of the genes fully and another 5 partly. Thus SDip allows the identification and characterization of paralog-specific phased sequence and structural variant calls from paralogues.

**Fig. 3.**
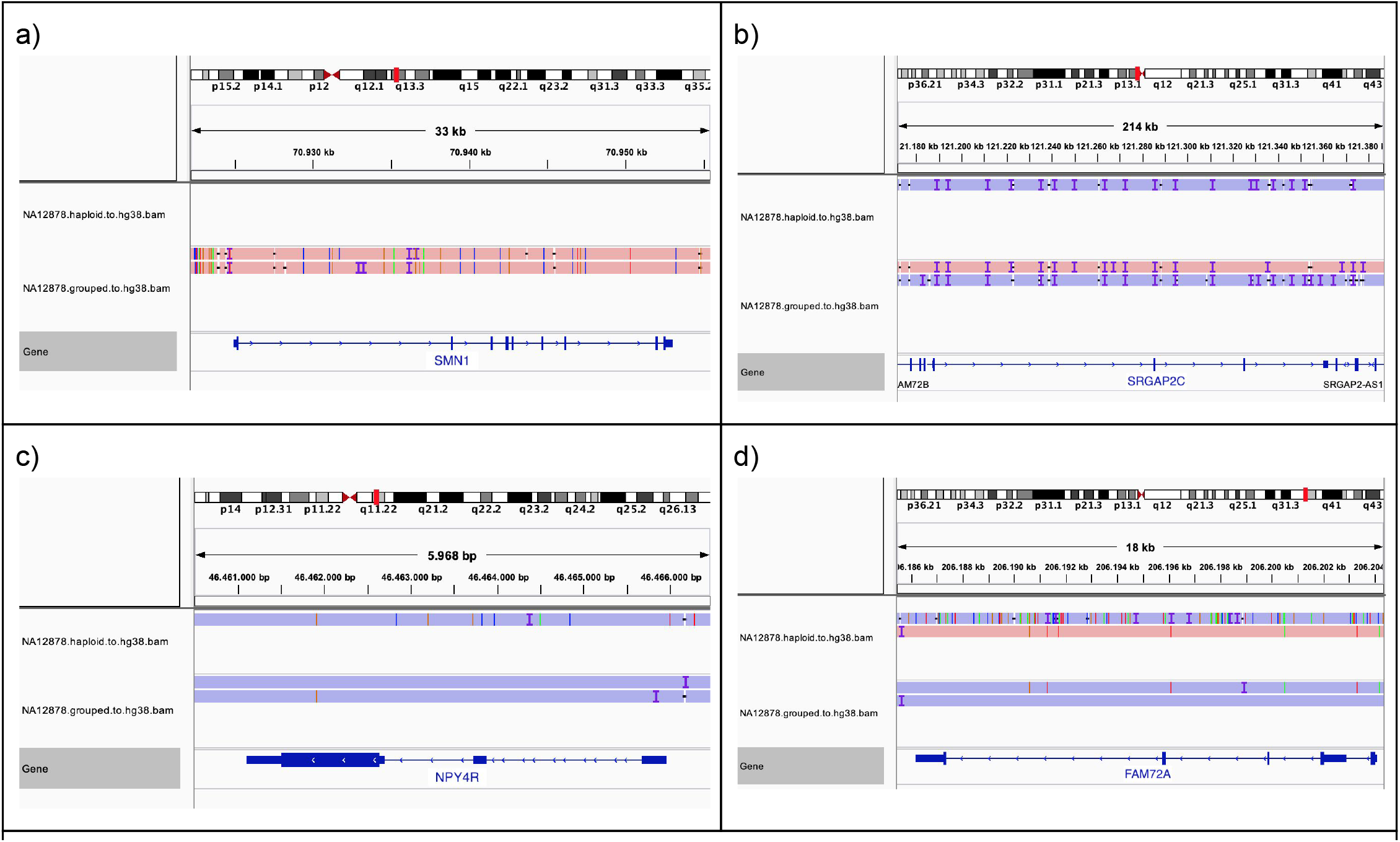
SDip contigs in biologically relevant genomic regions. SDA contigs (top panel) and SDip diploid contigs (bottom panel) in genes a) SMN1, b) SRGAP2C, c) NPY4R and d) FAM72A

## Discussion

Sequence-resolving of duplicated regions and their accurate variant calling in the human genome has been a challenging task due to the difficulty of distinguishing alleles and paralogues copies. In this study, we presented a new graph-based method that uses SNP information for distinguishing alleles and paralogues from overlapping reads to spell out phased contigs. These phased contigs are used for structural variant calling in these duplicated regions. Specifically, we tailored our method to leverage allelic and duplication-specific information from long accurate PacBio HiFi data. From our experiments on three genomes, we have demonstrated that we can fully assemble 86% of collapsed duplicated regions with continuity of more than 79 Kb and base level accuracy of more than 28 against comparison to BACs.

Our approach has some limitations. First, we restricted our analysis to segmental duplication region selection from the mixed haplotype assembly. We want to explore region selection from reads mapped to known segmental duplication genes in the reference genome or the phased assembly. The other potential region selection approach is to use whole-genome all-vs-vs overlapping reads. Second, we don’t assemble regions that result in graphs with cycles. One potential solution to resolve cyclic graphs is by incorporating other data types such as optical maps from bionano.

This approach is relevant for assembling polyploids, irrespective of its ploidy nature. One bottleneck is all-vs-all base level alignment for whole genomes that will be explored in the future. Nevertheless, our improved phased contigs and variant calls set up a milestone in duplicated regions. Our method enables fully phased contigs and structural variants in the duplicated regions for the first time. These advancements would open up the possibility to assemble phased contigs and variant calls in highly repetitive regions such as centromeres and ribosomal DNAs. The methodology behind SDip is generalized enough to be integrated into upcoming assemblers for phased whole-genome assembly, by applying this strategy to every read to distinguish reads from different haplotypes and repeats in complex regions.

## Methods

### Human genome assemblies

We considered three human genome assemblies derived from haploid (CHM13) and diploid genomes (NA12878 and HG002). All three genome assemblies were generated using high-coverage PacBio HiFi data. For CHM13, the assembly was generated using Canu on 24x PacBio HiFi data ^9^. The assemblies for HG002 and NA12878 were generated using Peregrine ^21^ on ~30x PacBio HiFi and then scaffolded using HiRise/3d-dna on ~30x Hi-C data ^22^.

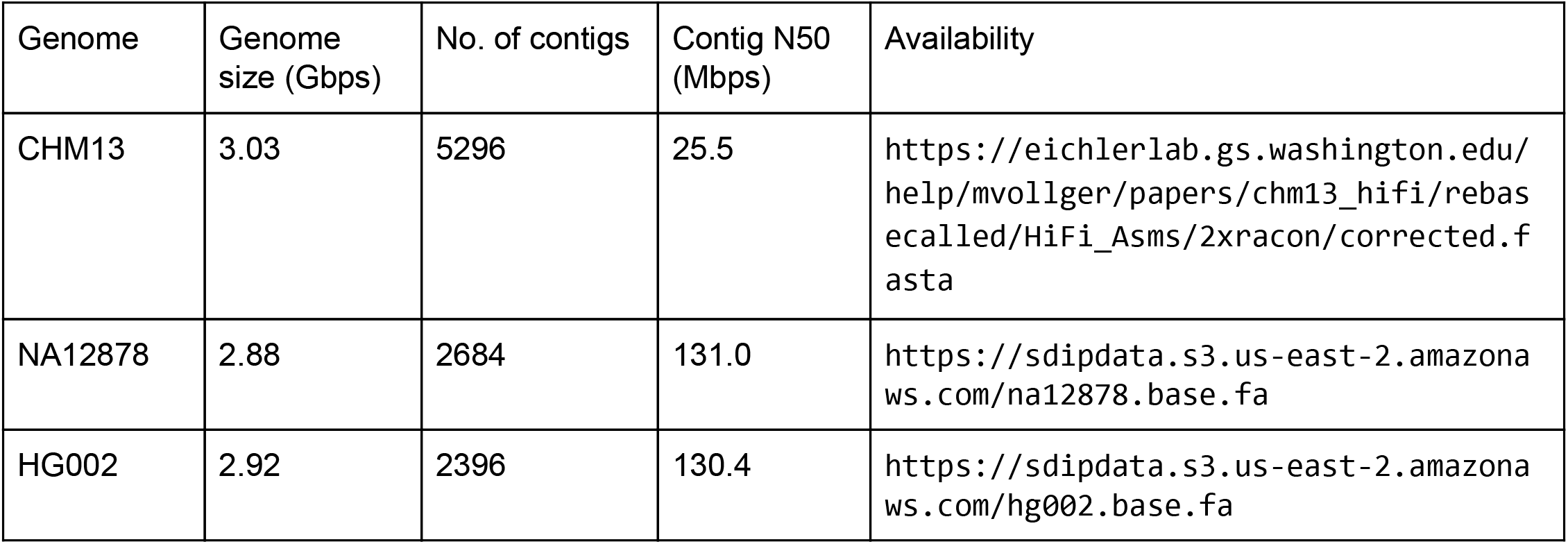

### BAC sequences

To validate our contigs, we used two sets of sequenced and assembled large-insert BAC clones for NA12878 and CHM13. Both datasets are publically available at NCBI.

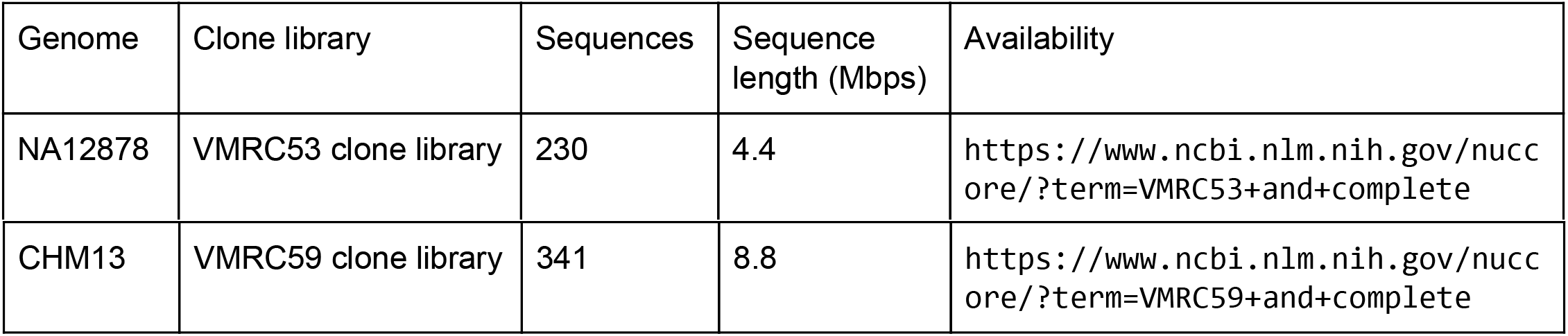

### SD detection and assembly

We used the tool SDA by Vollger et al. for detecting collapsed SD regions based on alignment depth of PacBio HiFi reads on the base assembly. Additionally, however, we aligned each collapsed sequence back to the base assembly with minimap2 ^23^ (`minimap2 -ax asm10 --secondary=yes -N 100 -p 0.80`) to find similar regions. We then recruited and combined PacBio HiFi reads from the collapsed region and all similar regions.

On the recruited reads from every region, we performed all-vs-all alignment using minimap2 ^23^ (`minimap2 -c -x asm20 -DP --no-long-join --cs -n500`). From the all-vs-all alignments, we considered the pileup for every read and detected true heterozygous SNVs including repeat or haplotype variation under the condition that allelic depth fraction > 0.11 and < 0.89. Based on these heterozygous informative sites, we decide if the reads from the pileup are from the same haplotype and repeat copy as the target read. While doing this, we follow a greedy heuristic to make an initial set of clusters based on overlaps of more than 4000 bp length and >15% of total number of variants in overlaps. Once these initial clusters are formed, we include another remaining set of reads into these clusters such that the hamming distance within clusters is minimized. We repeat the whole procedure for every read. To construct the overlap graph, the reads that are put into the same cluster as the target read have edges in the overlap graph. Additionally, however the reads with no variation within their overlaps, but have good overlaps of at least 4000 bp, there is an edge between them. This results in an haplotype- and repeat-aware overlap graph. Subsequently, we applied a transitive reduction of edges.

Next, we applied multiple graph cleaning steps (Figure S1). First, we pruned tips, i.e. linear chains of five or less nodes branching off from another chain. Secondly, we simplified bubbles, i.e. patterns in the graph where a start and an end node are connected by multiple node paths. To detect bubbles, we followed the algorithm by Onodera et al. and subsequently removed all but one path through the bubble ^24^. Finally, we removed cross-linking paths from the graph. Such erroneous paths between two nodes with a degree larger than 2 often link two branches of a large bubble. By removing all cross-links with a length of up to 4 nodes, we were able to clean up the graph considerably.

To this cleaned graph, we aligned ultralong nanopore reads from the same region. We specifically utilized alignment paths that connect two ends in the graph and thus represent a complete walk through the entire graph. To produce haplotype sequences, we spelled out a first set of contigs along the nanopore alignment paths. Additionally, we computed the minimum-cost minimum path cover for the graph, i.e. the smallest set of paths that traverse every node in the graph at least once. For every node, we computed the count of minimum path cover paths it is contained in using the tool mc-mpc-solver^25^. After subtracting the first nanopore-derived set of contigs from the node counts, we spelled out a second set of contigs through nodes with a count > 0 until all node counts are 0 and all nodes are contained in at least one contig.

Next, a haplotype-aware polishing of the contigs was performed. Note that every contig represents one path of PacBio HiFi reads in the overlap graph. Therefore, we recruited all reads associated with a contig and aligned them to the contig with minimap2 (`minimap2 -ax map-pb`). Then, we produced a pileup using bcftools mpileup (`bcftools mpileup -Q0 -o20 -e10 -Ov -f`), filtered for SNPs (`bcftools view -m2 -M3 -v snps -i ’DP > 3 & (I16[2] + I16[3]) / (I16[0] + I16[1]) > 1’ -Oz`) and Indels (`bcftools view -m2 -M2 -v indels -i ’DP > 3 & (I16[2] + I16[3]) / (I16[0] + I16[1]) > 1 & IMF>0.5’ -Oz`) using bcftools view and lastly produced a consensus with bcftools consensus (default parameters).

In a final step, we matched contigs that likely represent the two chromosomal haplotypes of the same paralogue. To this end, we performed all-vs-all alignment of the contigs from a given region using minimap2 (`minimap2 -x asm20 -Y -a --eqx`). Initially, we removed contigs with a percent identity of more than 99.95% to another contig. Then, we applied a flexible identity threshold to separate paralogous contigs with an identity lower than the threshold and allelic contigs with an identity higher than the threshold. For every region, the threshold was chosen that best separated the contigs into pairs of contigs with an identity higher than the threshold within each pair and an identity lower than the threshold between pairs. Even with the optimal threshold, not all contigs could be paired producing two sets of contigs: diploid contig pairs and the remaining haploid contigs. For the haploid genome CHM13, no matching was performed yielding only haploid contigs.

### Contig validation using BAC sequences

We validated contigs from SDip and SDA for NA12878 and CHM13 with publicly available BAC sequences. To estimate the QV of the assembled sequences, we aligned the BACs to the contigs with minimap2 (`minimap2 -I 8G --secondary=no -a --eqx -Y -x asm20 -m 10000 -z 10000,50 -r 50000 --end-bonus=100 -O 5,56 -E 4,1 -B 5`). ….

## Data availability

All assemblies, contigs and SV calls are available using `aws s3 sync --no-sign-request s3://sdipdata/`.

## Code

The code of the SDip pipeline and all evaluation scripts are available at: https://github.com/shilpagarg/sdip.git

## Author contributions

S.G. and H.L. conceived the study. H.L. proposed the idea of a ploidy agnostic algorithm. S.G. and D.H. implemented the algorithm. D.H. and S.G. implemented the pipeline and performed experiments. D.H., H.L., M.V., G.M.C. and S.G. analyzed and evaluated the diploid contigs. D.H., H.L. and S.G. wrote the manuscript. All authors helped to revise the manuscript.

---

**Fig. S1.**
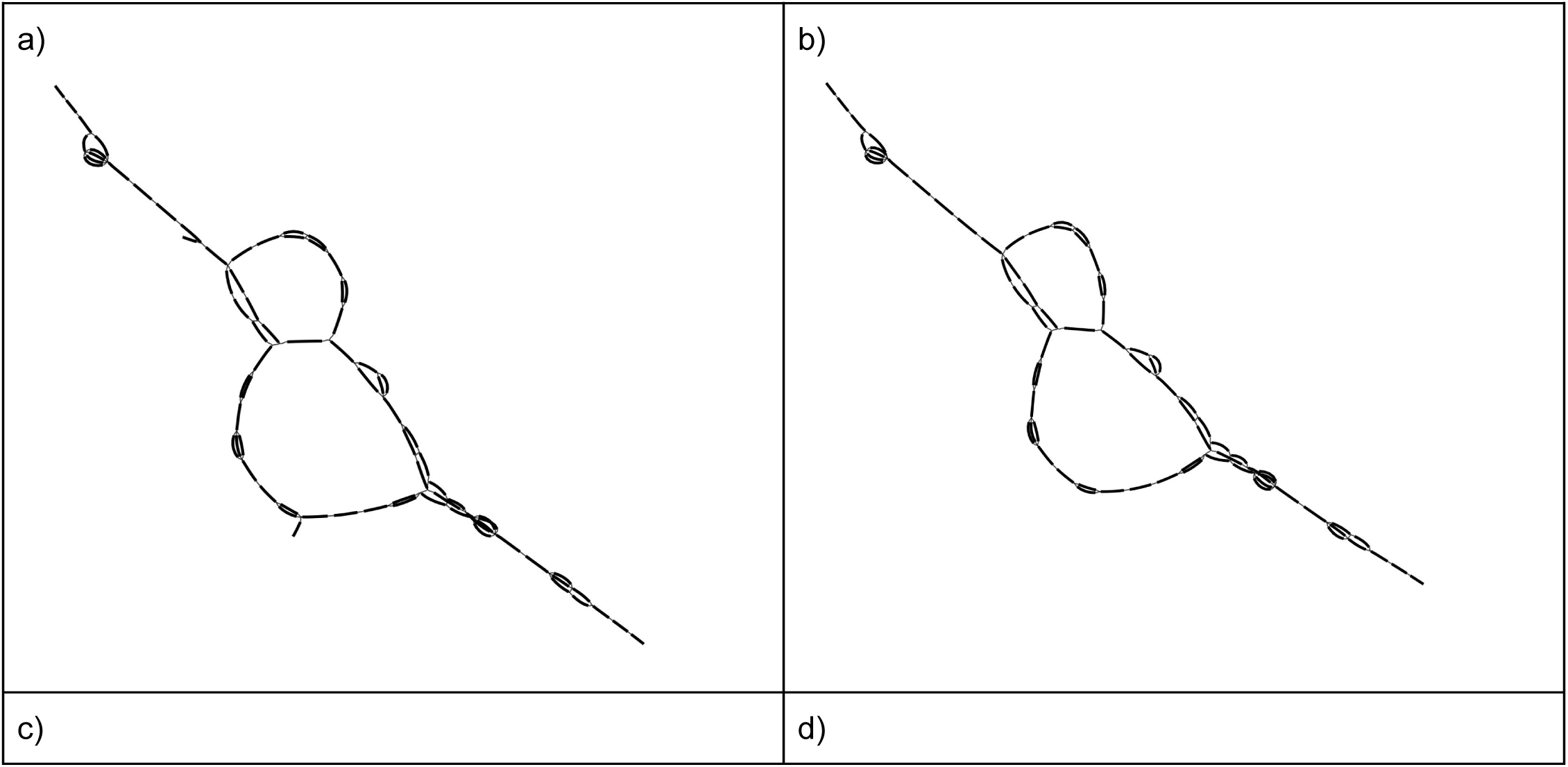
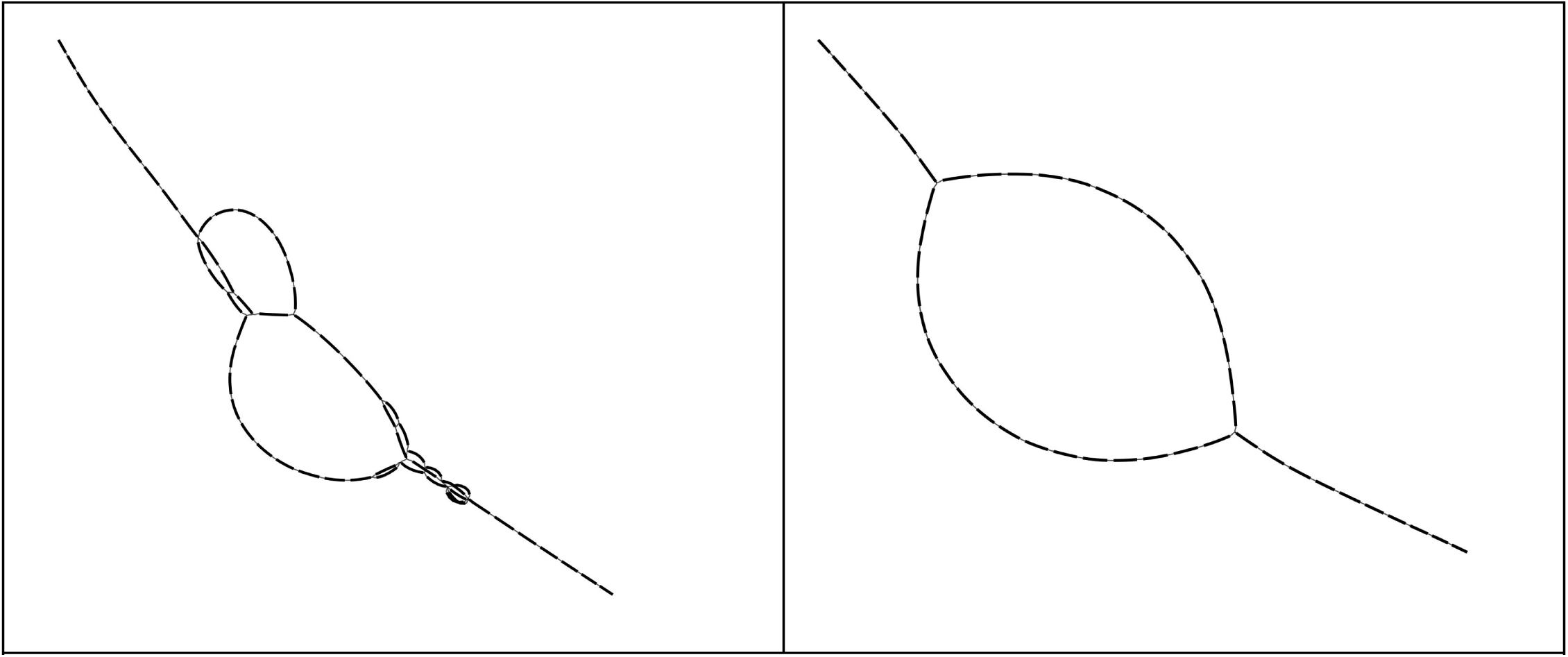
Pruning steps on overlap graph. a) Initial overlap graph with superfluous nodes and edges, b) short tips are removed, c) bubble simplification transforms short bubbles into linear node chains, d) removing edges between high-degree nodes resolves complex parts of the graph.

